# Spo13 prevents premature cohesin cleavage during meiosis

**DOI:** 10.1101/488312

**Authors:** Stefan Galander, Rachael E. Barton, David A. Kelly, Adèle L. Marston

## Abstract

Meiosis produces gametes through two successive nuclear divisions, meiosis I and meiosis II. In contrast to mitosis and meiosis II, where sister chromatids are segregated, during meiosis I, homologous chromosomes are segregated. This requires the monopolar attachment of sister kinetochores and the loss of cohesion from chromosome arms, but not centromeres, during meiosis I. The establishment of both sister kinetochore mono-orientation and cohesion protection rely on the budding yeast meiosis I-specific Spo13 protein, the functional homolog of fission yeast Moa1 and mouse MEIKIN. Here we investigate the effects of loss of *SPO13* on cohesion during meiosis I. Unlike wild type, cells lacking *SPO13* fail to maintain the meiosis-specific cohesin subunit, Rec8, at centromeres and segregate sister chromatids to opposite poles during anaphase I. We show that the cohesin-destabilizing factor, Wpl1, is not primarily responsible for the loss of cohesion during meiosis I. Instead, premature loss of centromeric cohesin during anaphase I in *spo13Δ* cells relies on separase-dependent cohesin cleavage. Further, cohesin loss in *spo13Δ* anaphase I cells is blocked by forcibly tethering the regulatory subunit of protein phosphatase 2A, Rts1, to Rec8. Our findings indicate that separase-dependent cleavage of phosphorylated Rec8 causes premature cohesin loss in *spo13Δ* cells.

## Introduction

Sexual reproduction relies on a cell division programme called meiosis. In humans, this is highly error-prone and may give rise to infertility, miscarriage or chromosomal abnormalities such as Down syndrome (reviewed in Hassold and Hunt 2001). Meiosis consists of two consecutive divisions where homologous chromosome segregation in meiosis I is followed by mitosis-like sister chromatid segregation in meiosis II. Homologue segregation requires a number of adaptations to the chromosome segregation machinery (Marston and Amon 2004), including recombination of homologues, mono-orientation of sister kinetochores and the protection of pericentromeric cohesin in meiosis I.

Cohesin is a multi-subunit protein complex made up of the core subunits Smc1, Smc3 and the kleisin α-Scc1 (Michaelis et al. 1997; Losada et al. 1998) as well as the accessory subunits Scc3 (Tóth et al. 1999) and Pds5 (Hartman et al. 2000; Panizza et al. 2000). In mitosis, cohesin resists the spindle forces that pull sister chromatids towards opposite poles, likely by topologically linking sister chromatids (Haering et al. 2002; Gruber et al. 2003). Upon successful bi-orientation, securin (Pds1 in yeast) is ubiquitinated and destroyed by the proteasome, freeing separase (Esp1), which proteolytically cleaves Scc1 and thereby allows chromosome segregation.

Meiotic cohesin contains an alternative kleisin called Rec8 (Buonomo et al. 2000; Watanabe and Nurse 1999). Rec8 supports a number of meiosis-specific functions of cohesin, particularly during recombination. Rec8 cleavage is dependent on its prior phosphorylation by three kinases – casein kinase 1δ (Hrr25), Dbf4-dependent kinase (DDK) Cdc7 (Katis et al. 2010) and Polo kinase (Cdc5) (Brar et al. 2006). However, it is currently unclear how these kinases contribute to cohesin removal with the role of Cdc5 in cohesin cleavage coming under particular scrutiny (Brar et al. 2006; Katis et al. 2010; Attner et al. 2013; Argüello-Miranda et al. 2017). While cohesin phosphorylation occurs along the length of the chromosome, the pericentromeric adapter protein shugoshin (Sgo1) binds protein phosphatase 2A (PP2A) to dephosphorylate Rec8 in the pericentromere and prevent its cleavage (Kitajima et al. 2004; Marston et al. 2004; Katis et al. 2004a; Kitajima et al. 2006; Tang et al. 2006; Riedel et al. 2006; Lee et al. 2008). In meiosis II, Rec8 becomes deprotected by the action of Hrr25, which is thought to initiate Sgo1 degradation and phosphorylate Rec8 for cleavage (Argüello-Miranda et al. 2017; Jonak et al. 2017).

In mammalian and *Drosophila* mitosis, cohesin is also removed in two steps. First, during prophase, Wapl opens the cohesin ring at the Smc3-Scc1 interface to trigger separase- and cleavage-independent cohesin removal (Waizenegger et al. 2000; Warren et al. 2000; Sumara et al. 2000; Buheitel and Stemmann 2013). A subset of cohesin is resistant to Wapl, owing to its acetylation and association with sororin (Rankin et al. 2005; Schmitz et al. 2007; Ben-Shahar et al. 2008; Unal et al. 2008; Lafont et al. 2010; Nishiyama et al. 2010). Notably, pericentromeric cohesin is shielded from Wapl during mammalian mitosis by Sgo1-PP2A which associates with, and dephosphorylates, both cohesin and sororin to prevent cohesin ring opening (Kitajima et al. 2006). Second, upon sister kinetochore bi-orientation, Sgo1 relocalises to the pericentromeric chromatin, and separase-dependent cohesin cleavage triggers anaphase onset (Liu et al. 2013b; 2013a).

While previous research has identified key mechanisms governing cohesin protection, a number of additional proteins have been implicated in this process, but their roles remain unclear. Amongst them is the meiosis I-specific Spo13 (Wang et al. 1987). Cells without *SPO13* only undergo a single meiotic division and show a variety of meiotic defects, including failure to mono-orient sister kinetochores in meiosis I and inability to protect cohesin (Klapholz and Esposito 1980; Shonn et al. 2002; Lee et al. 2004; Katis et al. 2004b). Spo13 is thought to have functional orthologs in both fission yeast (Moa1) and mouse (MEIKIN) (Kim et al. 2014). The unifying feature of these proteins is their interaction with Polo kinases, whose kinetochore recruitment by Moa1 and MEIKIN has been proposed to enable monoorientation and cohesin protection (Matos et al. 2008; Kim et al. 2014; Miyazaki et al. 2017).

The exact role of Spo13 in cohesin protection is currently unclear. Although it has been implicated in ensuring the proper pericentromeric localisation of Sgo1 (Kiburz et al. 2005), other studies have found no difference in chromosomally associated Sgo1 (Lee et al. 2004). In fact, it has been suggested that *spo13Δ* cells might retain residual pericenteromic cohesion in meiosis I (Katis et al. 2004b).

Here, we take a live cell imaging approach to re-evaluate the importance of Spo13 for cohesin protection. We show that both cohesin and sister chromatid cohesion are lost upon anaphase I onset in *spo13Δ* cells. Furthermore, cohesin removal results from separase-mediated cleavage rather than removal by the prophase pathway. We also provide evidence that cohesin phosphorylation is required for loss of cohesion in *spo13Δ* cells.

## Results and Discussion

### Pericentromeric cohesin is prematurely lost in *spo13Δ* cells

Previous analyses of fixed cells found that centromeric Rec8 is undetectable or greatly diminished in *spo13Δ* anaphase I cells (Klein et al. 1999; Katis et al. 2004b; Lee et al. 2004). Moreover, inactivation of *SPO13* allows *mam1Δ* cells (which lack sister kinetochore mono-orientation) to segregate sister chromatids during anaphase I (Katis et al. 2004b; Lee et al. 2004). Together, these findings provide evidence that centromeric cohesion is impaired in *spo13Δ* cells. However, it has been argued that residual centromeric cohesin persists after securin destruction in *spo13Δ* cells and prevents timely spindle elongation (Katis et al. 2004b). To clarify the importance of Spo13 in centromeric cohesion, we used live cell imaging of cells progressing through meiosis. We scored the percentage of cells where cohesin (Rec8-GFP) was retained at the pericentromere in anaphase I, as indicated by co-localisation with Mtw1 (Figs. 1A and 1B). To ensure that observed effects in *spo13Δ* cells were not a consequence of mono-orientation loss, which partially impacts cohesion (Nerusheva et al. 2014), we simultaneously imaged *mam1Δ* cells for comparison. Quantification of pericentromeric Rec8 (Fig. 1C) showed that, strikingly, deletion of *SPO13* leads to complete loss of cohesin in anaphase I. This is not due to impaired cohesin loading in early meiosis, since prophase I-arrested *spo13Δ* cells have similar levels of Rec8 on centromeres compared to wild type (Fig. 1D). We conclude that Spo13 is required for the retention of pericentromeric cohesin in anaphase I.

**Figure 1.**
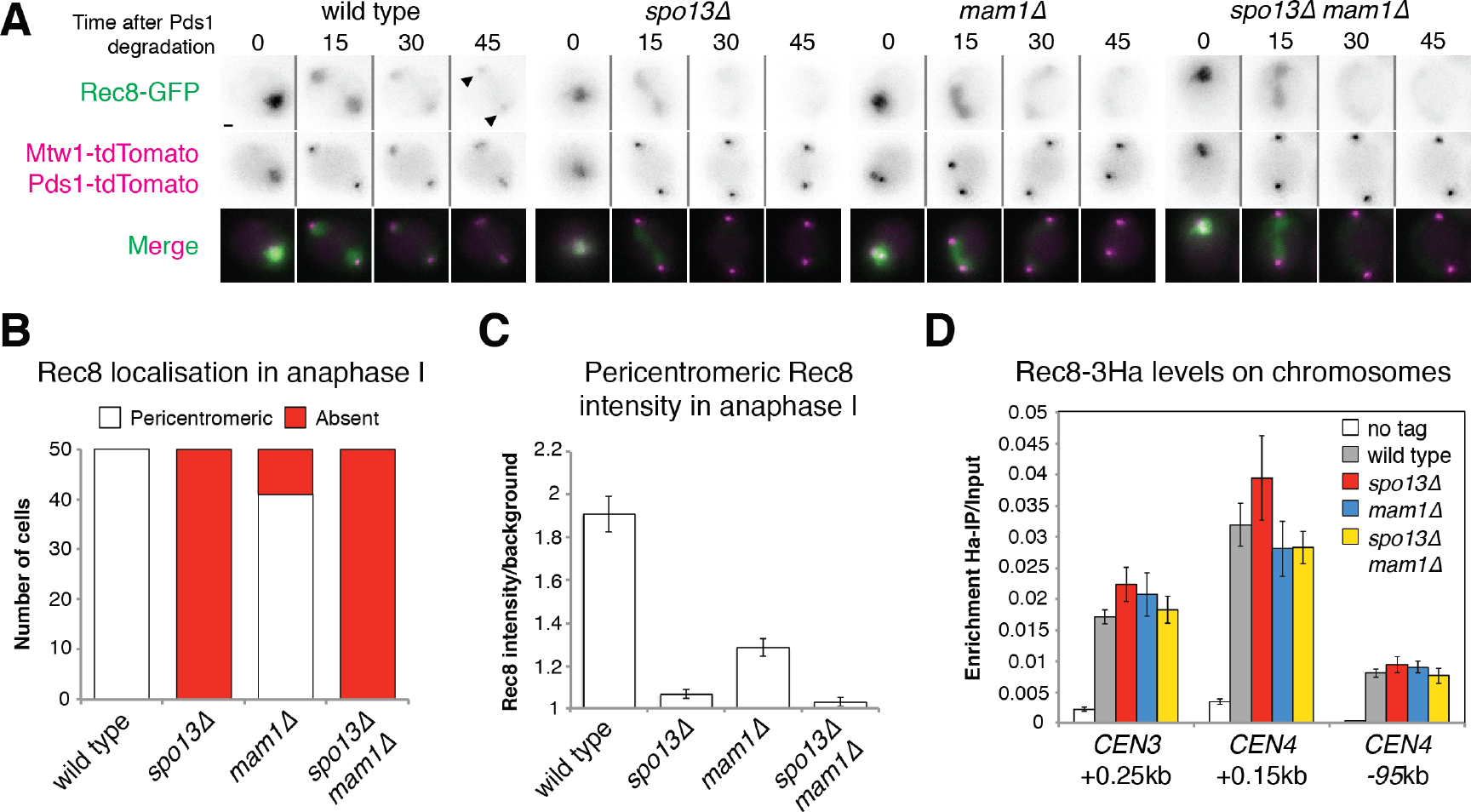
Cohesin and sister chromatid cohesion are lost in the absence of *SPO13*. (A) Representative images of Rec8-GFP, Mtw1-tdTomato and Pds1-tdTomato in live sporulating wild type (AM13716), *spo13Δ* (AM15133), *mam1Δ* (AM15134) and *spo13Δ mam1Δ* (AM15135) cells. Scale bars represent 1μm. Arrows indicate pericentromeric cohesin. (B) The number of cells with pericentromeric Rec8-GFP in anaphase I is shown after scoring 50 cells from (A). (C) Rec8-GFP intensity was measured for 50 cells from (A) in the area occupied by the tdTomato-labeled kinetochore protein Mtw1. (D) Rec8 loading is unaffected by deletion of *SPO13*. Rec8-3HA association with the indicated sites was measured in prophase I in wild type (AM4015), *spo13Δ* (AM15343), *mam1Δ* (AM15342) and *spo13Δ mam1Δ* (AM15344) cells carrying *ndt80Δ* and a no tag control (AM11633). Cells were arrested in prophase by harvesting 5 h after resuspension in sporulation medium and anti-Ha ChIP-qPCR performed.

### *spo13Δ* cells prematurely segregate sister chromatids

To assess sister chromatid cohesion in *spo13Δ* cells, we labelled one copy of chromosome V near the centromere with an array of tet operators (*tetO*), expressed GFP-tagged TetR repressor (Michaelis et al. 1997) and imaged *CEN5*-GFP foci in live meiotic cells. Upon anaphase I entry (as judged by degradation of Pds1 (Salah and Nasmyth 2000)), three different phenotypes may be observed, depending on whether cells successfully mono-orient sister kinetochores and protect pericentromeric cohesin (Fig. 2A). In wild type cells, a single GFP focus segregates to one of the spindle poles (as marked by Spc42-tdTomato). Alternatively, in case of defective mono-orientation, split GFP foci stay in close proximity (<2μm) because sister chromatids together are cohered by pericentromeric cohesin. Lastly, in cells lacking both mono-orientation and sister chromatid cohesion, GFP dots split over a greater distance (>2μm). We subsequently scored the number of cells falling into either of these categories for each of the mutants analysed. This revealed that sister centromeres separate over large (>2 μm) distances in the half of *spo13Δ* anaphase I cells that bi-orient sister kinetochores (Fig. 2B), consistent with all cohesion being lost. Note that although pericentromeric cohesion loss during anaphase I can only be readily observed where it is accompanied by sister kinetochore bi-orientation, the loss of cohesion in all *spo13Δ* cells with bi-oriented kinetochores, the near-complete absence of Rec8, and the fact that deletion of *SPO13* permits efficient sister chromatid segregation in *mam1Δ* cells (Fig. 2B) (Lee et al. 2004; Katis et al. 2004b) together confirm that pericentromeric cohesion is non-functional in *spo13Δ* anaphase I cells.

**Figure 2.**
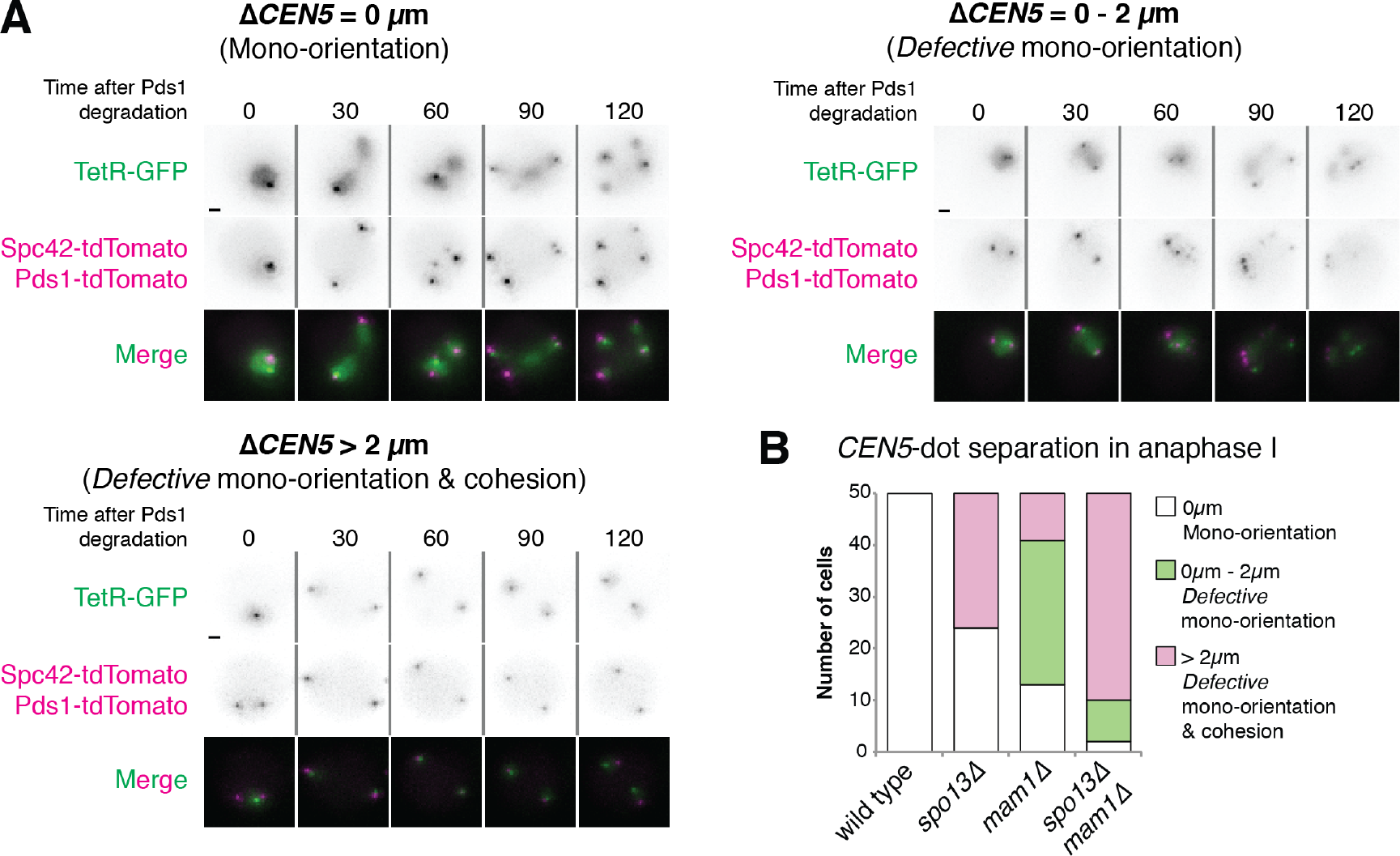
Deletion of *SPO13* permits sister chromosome segregation in anaphase I of *mam1Δ* mutants. (A) Assay for mono-orientation and cohesion defects using heterozygous centromeric fluorescent markers. Representative images are shown. Scale bars represent 1μm. Images for Δ*CEN5*=0μm, Δ*CEN5*=0-2μm and Δ*CEN5*>2μm were taken from wild type, *mam1Δ* and *spo13Δ* cells, respectively. (B) Frequency of *CEN5* distance categories is shown for the indicated genotypes after live cell imaging. Wild type (AM15190), *spo13Δ* (AM15118), *mam1Δ* (AM15119) and *spo13Δ mam1Δ* (AM15120) cells carrying *SPC42-tdTomato*, *Pds1-tdTomato* and heterozygous TetR-GFP dots at *CEN5*, were were sporulated for 2.5 h before imaging on a microfluidics plate.

### Sister chromatid cohesion is restored by preventing cohesin cleavage

A cleavage-independent, Rad61/Wpl1-dependent, cohesin removal pathway, similar to that which occurs in mammalian mitosis, operates during prophase I of budding yeast meiosis (Yu and Koshland 2005; Challa et al. 2016; 2018). We considered the possibility that cells lacking Spo13 lose cohesion, not due to its cleavage, but as a result of ectopic Rad61 activity. However, deletion of *RAD61* did not restore cohesion to *spo13Δ* cells (Fig. 3A), indicating that a failure to counteract cleavage-independent cohesin removal is not solely responsible for the cohesion defect of cells lacking Spo13.

**Figure 3.**
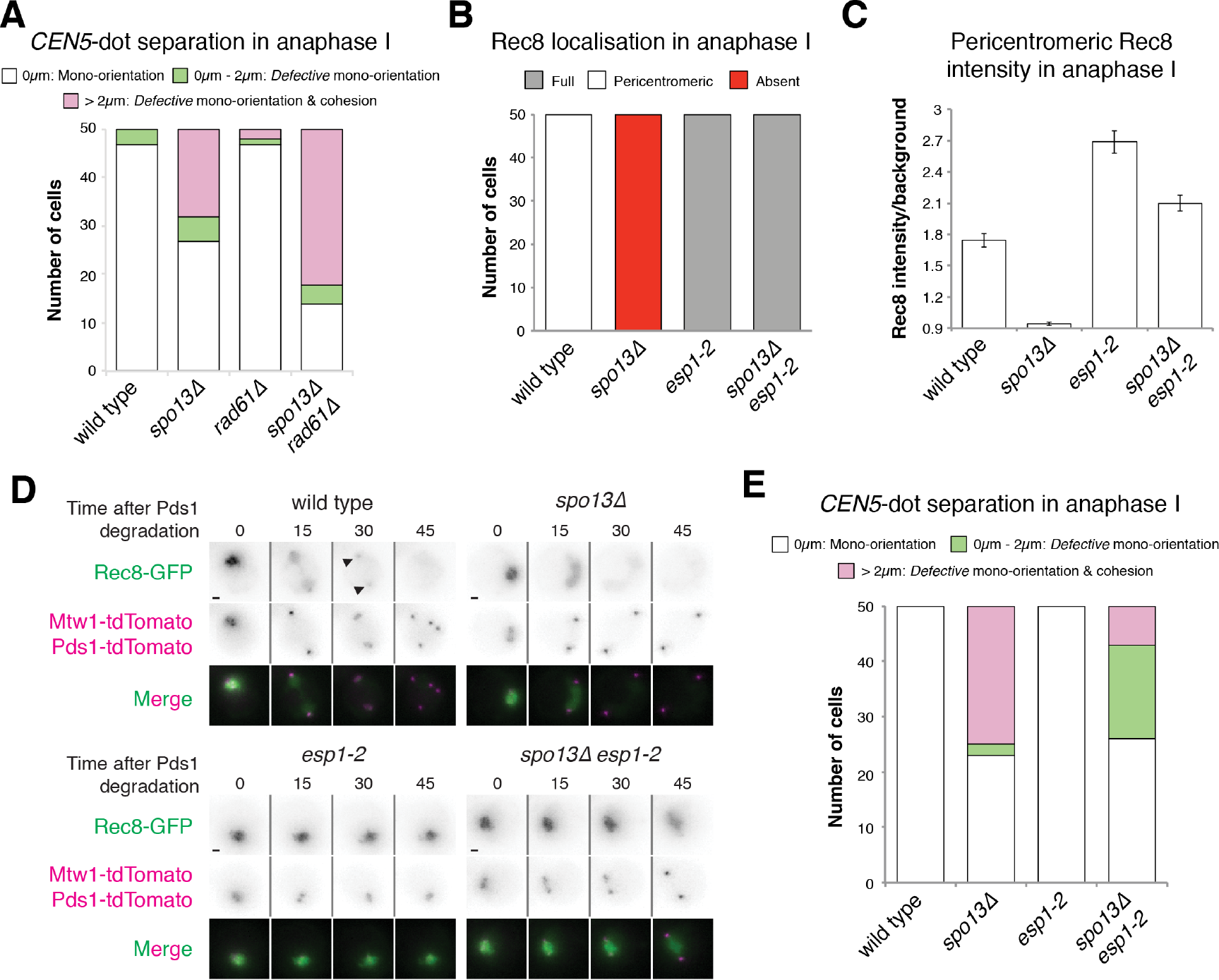
Cohesin protection in *spo13Δ* cells is rescued by inhibition of separase, but not by ablation of the prophase pathway. (A) Deletion of *RAD61/WPL1* does not rescue sister chromatid cohesion in *spo13Δ* cells. Categorisation of *CEN5-GFP* distances in wild type (AM15190), *spo13Δ* (AM20146), *rad61Δ* (AM21068) and *spo13Δ rad61Δ* (AM21358) cells carrying *SPC42-tdTomato*, *PDS1-tdTomato* and heterozygous TetR-GFP dots at *CEN5* was carried out as described in Fig. 2A. (B-D) Separase activity is required for Rec8 removal in *spo13Δ* mutants. Wild type (AM13716), *spo13Δ* (AM20033), *esp1-2* (AM20868) and *spo13Δ esp1-2* (AM21949) cells carrying *REC8-GFP, MTW1-tdTomato* and *PDS1-tdTomato* were resuspended in sporulation medium at 32°C and grown in flasks for 3h before transferring to a microfluidics plate and imaged at 32°C. (B) The number of cells with the indicated patterns of Rec8-GFP localisation in anaphase I was scored for 50 cells per strain. (C) The intensity of pericentromeric Rec8-GFP for the indicated genotypes is shown. The mean of the two maximum intensity values on a straight line connecting both kinetochores in anaphase I (within the first two time points after Pds1-tdTomato degradation) was measured for 50 cells. Error bars represent standard error. (D) Representative images are shown. Scale bars represent 1μm. Arrows indicate pericentromeric cohesin. (E) Inhibition of separase activity restores sister chromatid cohesion to *spo13Δ* mutants. Cohesion was assayed by categorisation of *CEN5*-GFP distances as described in Fig 1E. Strains used were wild type (AM15190), *spo13Δ* (AM20146), *esp1-2* (AM22498) and *spo13Δ esp1-2* (AM22499) cells carrying *SPC42-tdTomato*, *PDS1-tdTomato* and heterozygous TetR-GFP dots at *CEN5*.

Next, we asked whether cohesin cleavage is required for cohesion loss during anaphase I in *spo13Δ* cells. First, we inactivated Esp1 (separase), using the temperature-sensitive *esp1-2* mutant (Buonomo et al. 2000) and followed Rec8-GFP by live cell imaging (Figs. 3B-D). As expected, cohesin remained on chromosomes even after anaphase I onset in both in *esp1-2* and *esp1-2 spo13Δ* cells and, consequently, sister chromatid segregation was largely prevented (Fig. 3E).

Additionally, we prevented cohesin cleavage by mutating the separase cleavage site in Rec8 (Rec8-N) (Buonomo et al. 2000). We followed GFP-tagged versions of this Rec8 variant through meiosis in wild type and *spo13Δ* cells (Fig. 4A). Similar to *esp1-2* mutants, *rec8-N* prevents cleavage of cohesin along the length of the chromosome in *spo13Δ* cells (Fig. 4B) and pericentromeric cohesin intensity is drastically increased (Fig. 4C). Furthermore, we find that Rec8-N prevented the segregation of sister chromatids in *spo13Δ* mutants (Fig. 4D). We conclude that cohesin cleavage is required for sister chromatid segregation in *spo13Δ* cells.

**Figure 4.**
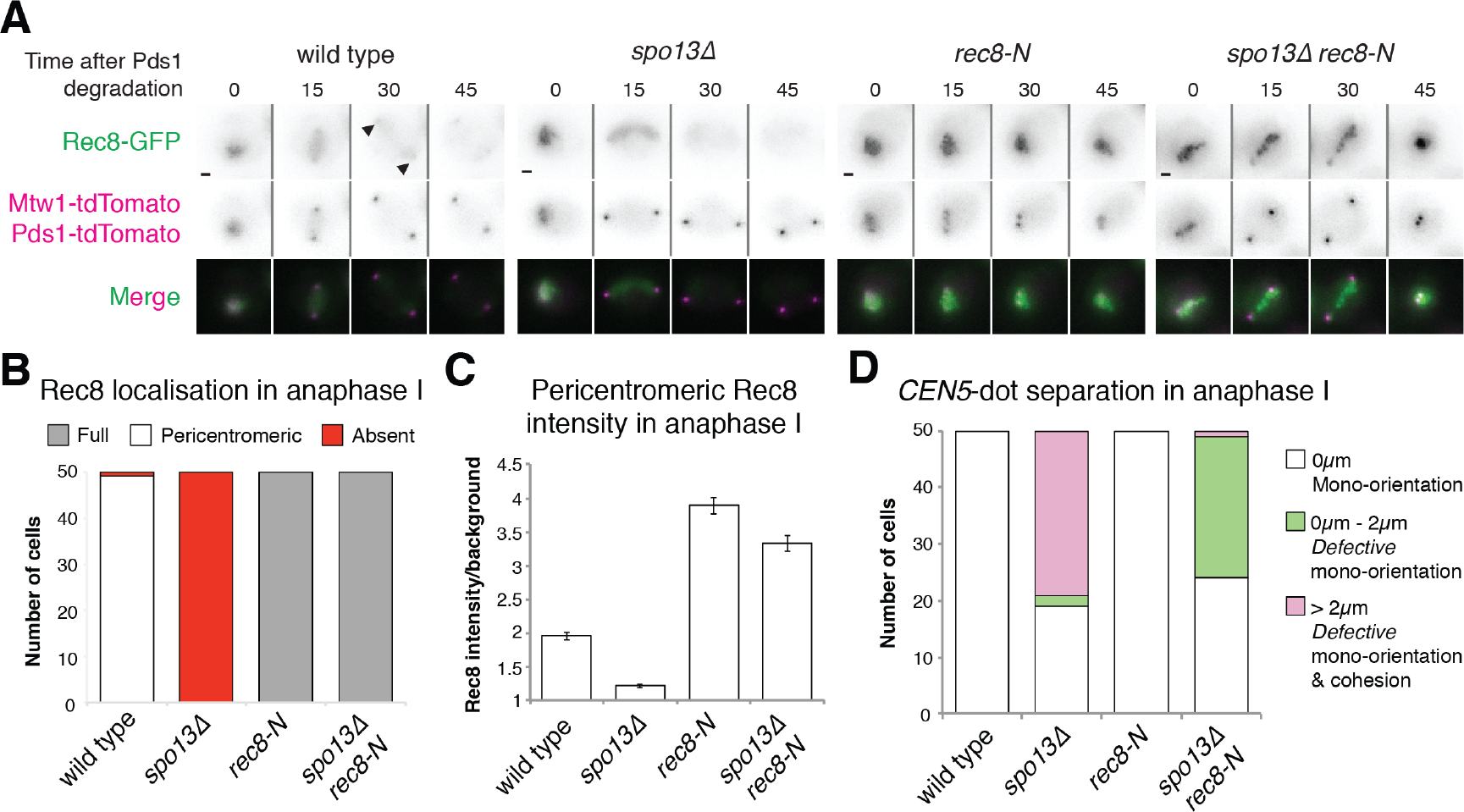
Cohesin cleavage is required for loss of sister chromatid cohesion in *spo13Δ* cells. (A-C) Non-cleavable Rec8 blocks efficient removal of cohesin in *spo13Δ* cells. (A) Representative images from movies of cells carrying Rec8-GFP, Mtw1-tdTomato, Pds1-tdTomato and with the indicated genotypes are shown. Scale bars represent 1μm. Arrows indicate pericentromeric Rec8-GFP. (B) Frequency of cells with the indicated pattern of Rec8-GFP localisation is shown for the indicated genotypes. (C) Rec8-GFP intensity was measured for the indicated genotypes as described in Fig. 3C. Error bars represent standard error. Strains used in (A-C) were *REC8-GFP* (AM22190), *REC8-GFP spo13Δ* (AM22191), *rec8-N-GFP* (AM22192) and *rec8-N-GFP spo13Δ* (AM22193) cells carrying *MTW1-tdTomato* and *PDS1-tdTomato*. (D) Non-cleavable Rec8 prevents sister chromatid segregation in *spo13Δ* mutants. Cohesion functionality was determined for the indicated genotypes by categorisation of *CEN5-GFP* distances as described for Fig. 2A. Strains were *REC8-3HA* (AM22346), *REC8-3HA spo13Δ* (AM22347), *rec8-N-3HA* (AM22348) and *rec8-N-3HA spo13Δ* (AM22349) and carried *SPC42-tdTomato*, *PDS1-tdTomato* and heterozygous TetR-GFP dots at *CEN5*.

### PP2A is functional in the absence of Spo13

Rec8 cleavage during wild type meiosis relies on its prior phosphorylation (Brar et al. 2006; Katis et al. 2010) which is reversed in the pericentromere by PP2A. We considered the possibility that PP2A function may be impaired in *spo13Δ* cells, rendering it unable to dephosphorylate, and therefore protect, cohesin. We asked whether tethering PP2A directly to cohesin could prevent Rec8 cleavage in the absence of Spo13. We fused GFP-binding protein (GBP), a nanobody specifically recognising GFP (Rothbauer et al. 2006), to the PP2A regulatory subunit Rts1 to irreversibly tether PP2A to GFP-tagged Rec8. This was sufficient to prevent cohesin removal, both in *pCLB2-SGO1* and *spo13Δ* cells (Figs. 5A-C). To further confirm the full functionality of Rts in *spo13Δ* cells, we utilised a separase biosensor (Yaakov et al. 2012) where a cleavable Rec8 moiety is fused to GFP and LacI, with the latter allowing targeting of the biosensor to a lacO array on chromosome arms (Fig. 6A). In wild type and *spo13Δ* cells, this biosensor appears as a single GFP focus in meiosis I until separase is activated in anaphase I, causing biosensor cleavage and GFP focus dispersal (Figs. 6B and C). Tethering of Rts1 to the biosensor, however, prevents biosensor cleavage (Figs. 6B and C). Therefore, our results indicate that PP2A is functional and capable of dephosphorylating cohesin in *spo13Δ* mutants.

**Figure 5.**
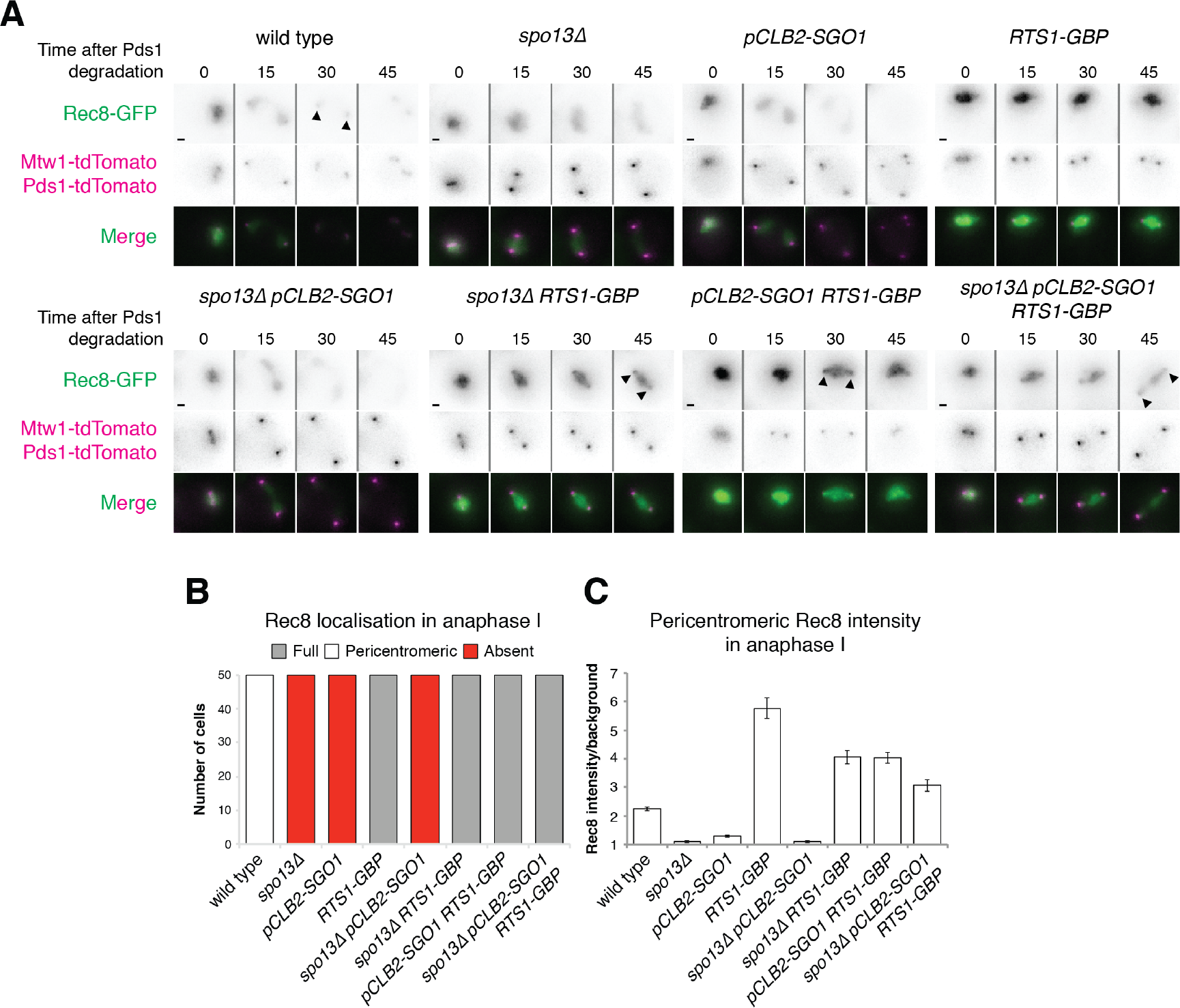
PP2A can prevent cohesin cleavage and sister chromatid segregation in *spo13Δ* cells. (A-C) Cohesin is retained on chromosomes when PP2A^Rts1^ is tethered to Rec8. (A) Representative images of Rec8-GFP, Mtw1-dtTomato and Pds1-tdTomato in wild type (AM13716), *spo13Δ* (AM20033), *pCLB2-SGO1* (AM21315), *RTS1-GBP* (AM21316), *spo13Δ pCLB2-SGO1* (AM21317), *spo13Δ RTS1-GBP* (AM21319), *pCLB2-SGO1 RTS1-GBP* (AM21318) and *spo13Δ pCLB2-SGO1 RTS1-GBP* (AM21320) cells undergoing meiosis. Scale bars represent 1μm. Arrows indicate pericentromeric cohesin. (B) The number of cells with pericentromeric cohesin in anaphase I was scored for 50 cells per strain. (C) Rec8-GFP intensity in anaphase I was measured as described in Fig. 2A.

**Figure 6.**
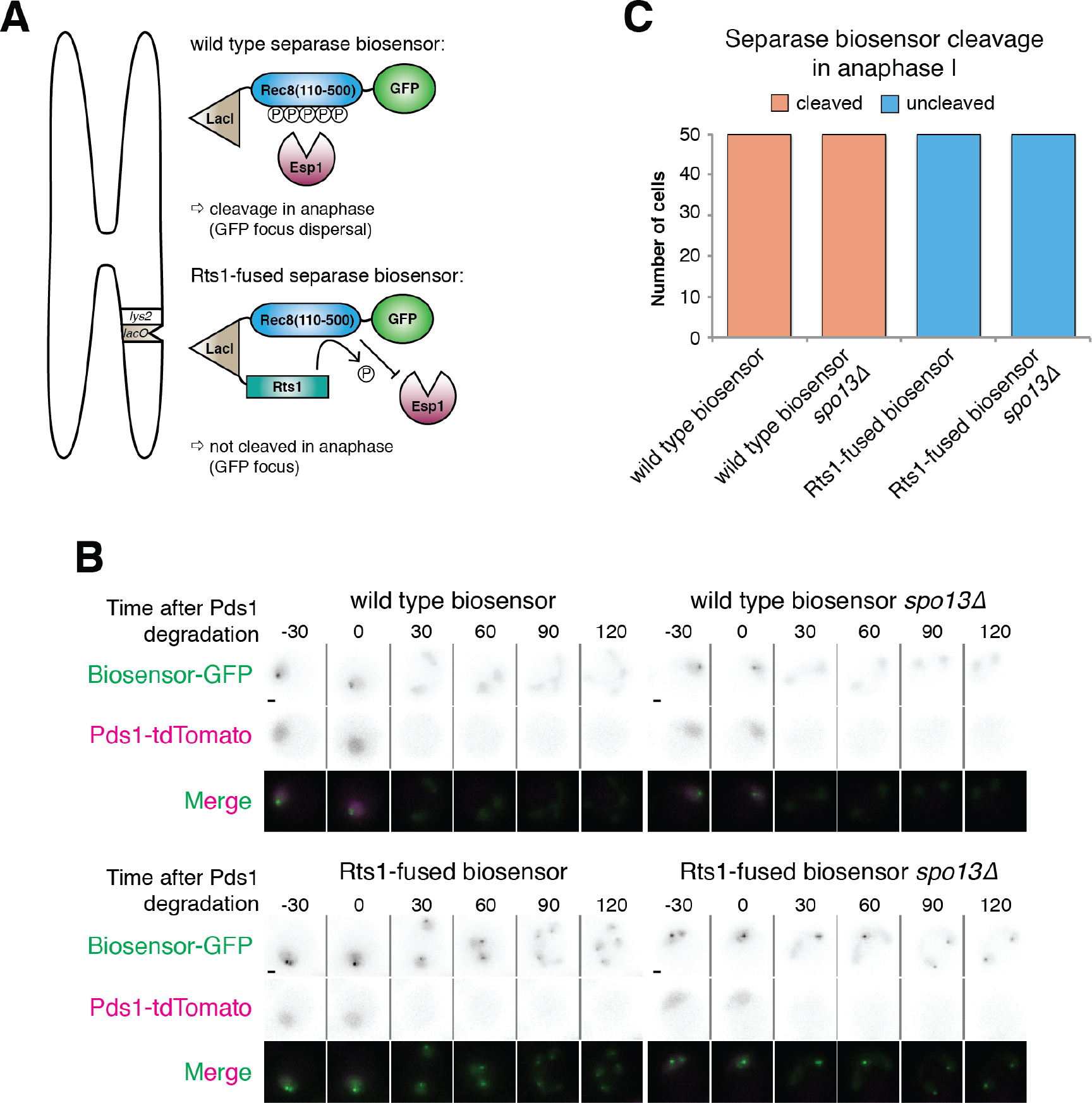
Fusion of Rts1 to a separase biosensor prevents its cleavage in both wild type and *spo13Δ* cells. (A) Schematic illustration of the separase biosensor and its Rts1 fusion. (B and C) Wild type (AM21557) and *spo13Δ* cells (AM21558) carrying a wild type separase biosensor (*pCUP1-GFP-REC8(110-500)-LacI*) or an Rts1 fused biosensor (*pCUP1-GFP-REC8(110-500)-LacI-RTS1;* wild type: AM21559, *spo13Δ:* AM21800) as well as *lys2::lacOx256* and *PDS1-tdTomato* were sporulated in the presence of 100nM CuSO_4_ for 2.5 h before imaging on a microfluidics plate. (B) Representative images are shown. Scale bars represent 1μm. (C) 50 cells per strain were scored for the presence of GFP foci (uncleaved biosensor) or diffuse GFP signal (cleaved biosensor) within 30 min (two time points) of Pds1 degradation.

## Conclusions

The successful protection of pericentromeric cohesin is a key modification to the meiotic chromosomes segregation machinery as it ensures the fidelity of chromosome segregation in meiosis II. Key players in regulating cohesin cleavage are known. The kinases Hrr25 and Cdc7 (and possibly Cdc5) phosphorylate cohesin along the length of the chromosome to promote its cleavage by separase (Brar et al. 2006; Katis et al. 2010; Attner et al. 2013), while pericentromeric Sgo1 recruits the phosphatase PP2A to dephosphorylate Rec8 and thereby protect it (Marston et al. 2004; Katis et al. 2004a; Kitajima et al. 2004; Riedel et al. 2006; Tang et al. 2006; Lee et al. 2008). However, a number of other factors, including the meiosis I-specific Spo13, are also required to retain pericentromic cohesin in anaphase I (Shonn et al. 2002; Lee et al. 2004; Katis et al. 2004b) but their functions are much less well understood. Our study demonstrates that pericentromeric cohesin is prematurely removed in *spo13Δ* cells in a manner that requires cohesin cleavage and phosphorylation. Future work should focus on elucidating how Spo13 elicits its protective function, and how this might be linked to its functions in both sister kinetochore mono-orientation and meiotic cell cycle control.

## Acknowledgements

We are very grateful to to Robin Allshire, Angelika Amon, Kevin Hardwick and David Morgan for sharing plasmids. We also thank members of the Marston lab for helpful discussion. This research was funded by Wellcome through PhD studentships to SG [096994] and REB [102316], as well as a Senior Research Fellowship to ALM [107827] and core funding awarded to the Wellcome Centre for Cell Biology [203149].

## Author contributions

SG and ALM conceived the study. SG performed all experiments except that shown in Fig. 3A, which was carried out by REB. DAK generated image analysis tools. SG prepared all figures. SG and ALM wrote the paper.

## Materials and Methods

### Yeast strains and plasmids

All strains are SK1-derivatives and are listed in Table S1. Plasmids generated in this study are listed in Table S2. Gene deletions, promoter replacements and gene tags were introduced using PCR-based methods (Knop et al. 1999; Longtine et al. 1998; Moqtaderi and Struhl 2008; Gauss et al. 2005). *pCLB2-CDC20* (Lee and Amon 2003), *REC8-GFP*, *Pds1-tdTomato* (Matos et al. 2008), *ndt80Δ* (Vincenten et al. 2015), *SPC42-tdTomato* (Fox et al. 2017), *REC8-3HA* (Klein et al. 1999), *CEN5*-GFP dots, *mam1Δ::TRP1* (Tóth et al. 2000) and *REC8-N* (Buonomo et al. 2000) were described previously. Separase biosensor constructs (Yaakov et al. 2012) were a kind gifts from David Morgan.

### Growth conditions

Cells were prepared for sporulation as described by Vincenten et al. (2015).

### Chromatin immunoprecipitation

ChIP-qPCR was performed as previously described (Vincenten et al. 2015), using mouse anti-Ha (12CA5, Roche). Primers for qPCR analysis are listed in Table S3.

### Live cell imaging

Live cell imaging was performed on a DeltaVision Elite system (Applied Precision) connected to an inverted Olympus IX-71 microscope with a 100x UPlanSApo NA 1.4 oil lens. Images were taken using a Photometrics Cascade II EMCCD camera. The Deltavision system was controlled using SoftWoRx software.

Cells were imaged at 30°C (unless stated) on an ONIX microfluidic perfusion platform by CellASIC. Cells were pre-grown in culture flasks for ~3 h before transfer to microfluidics plates. Imaging began about 30 min later with images being acquired every 15 min for 12-15 h. Seven z-stacks were acquired with 0.85μm spacing. Image panels were assembled using Image Pro Premier (Media Cybernetics). Images were analysed using ImageJ (National Institutes of Health). Final image assembly was carried out using Adobe Photoshop and Adobe Illustrator. Rec8-GFP intensities were measured a custom plugin for ImageJ. The plugin applied a Z projection to each colour channel and allowed the user to select a cell of interest. Kinetochores in the red channel were identified by Yen autothreshold and their XY central coordinates, mean intensity and area recorded. The coordinates were then used to measure mean intensity in the corresponding location in the green channel, equivalent to pericentromeric Rec8-GFP. In experiments where pericentromeric cohesin was likely to be found in between kinetochores (which is thought to occur in cells that bi-orient in meiosis I but retain cohesin), the XY coordinates in the red channel were used to generate a line profile between the 2 kinetochores in both colour channels over exactly the same pixels. The 2 brightest peaks in the line profile of the green channel were calculated to give the maximum intensity value for each. Rec8-GFP intensity was measured in this manner for Figs. 3C and 4C.

## Supplementary Information

**Table S1.**
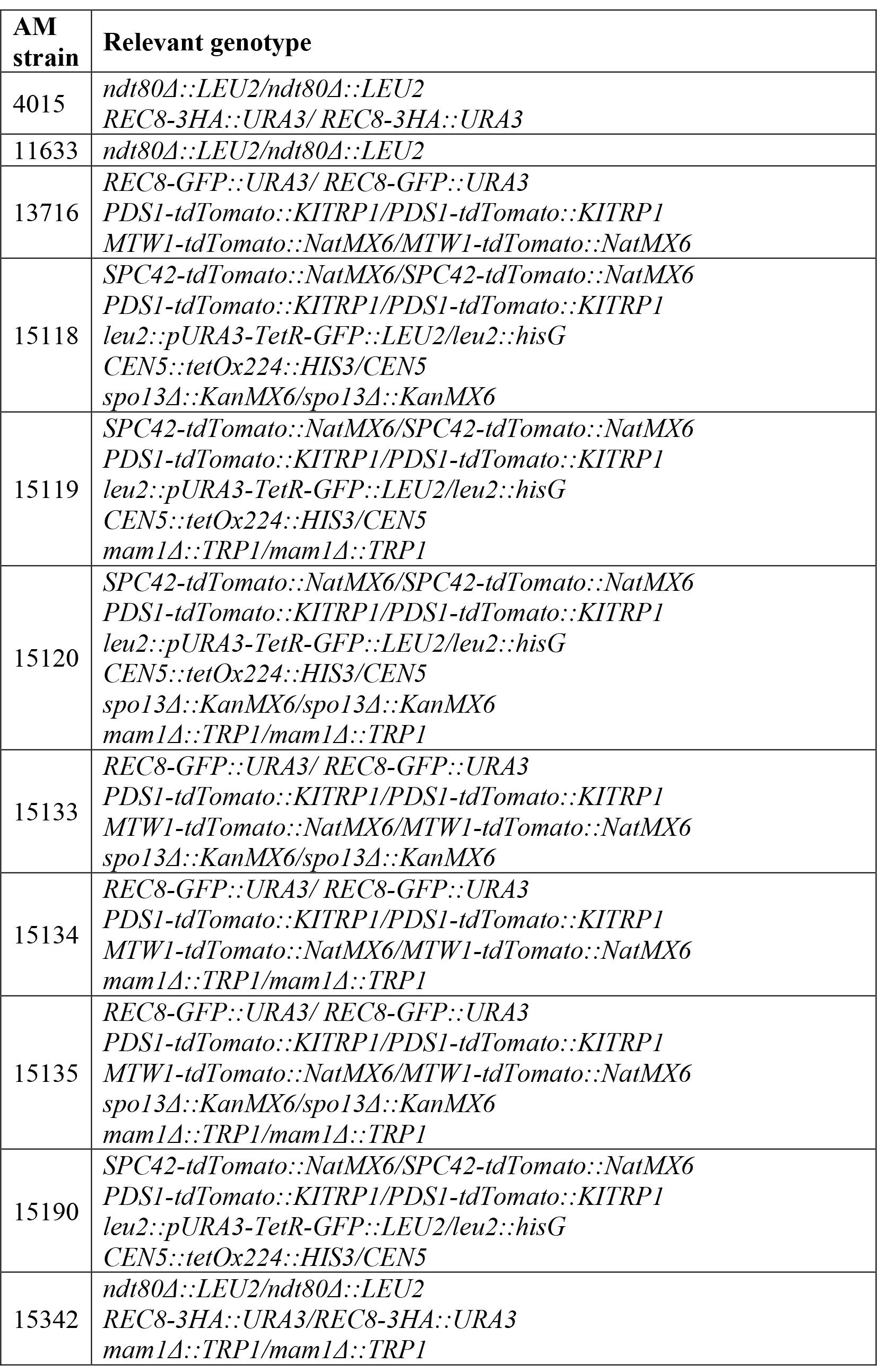
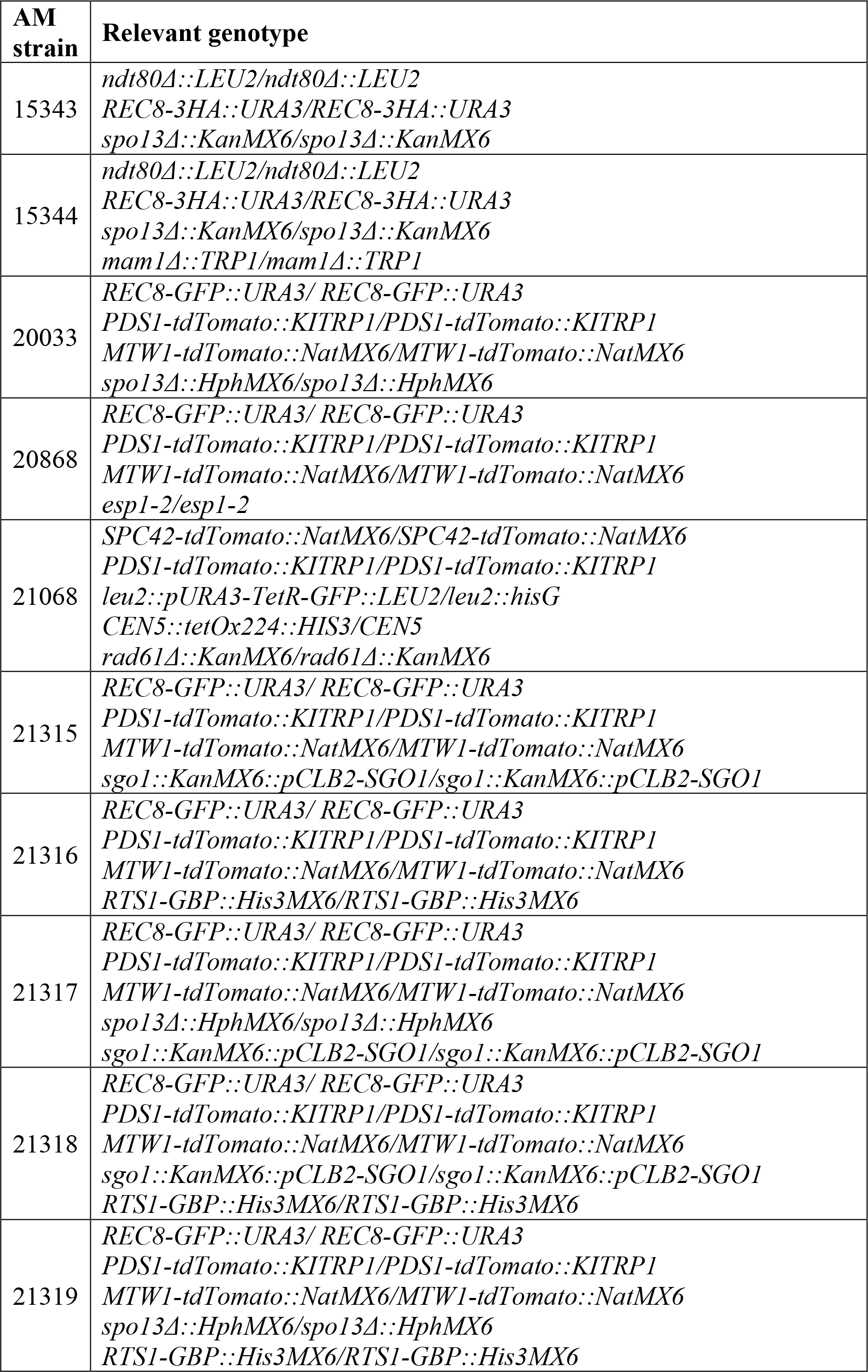
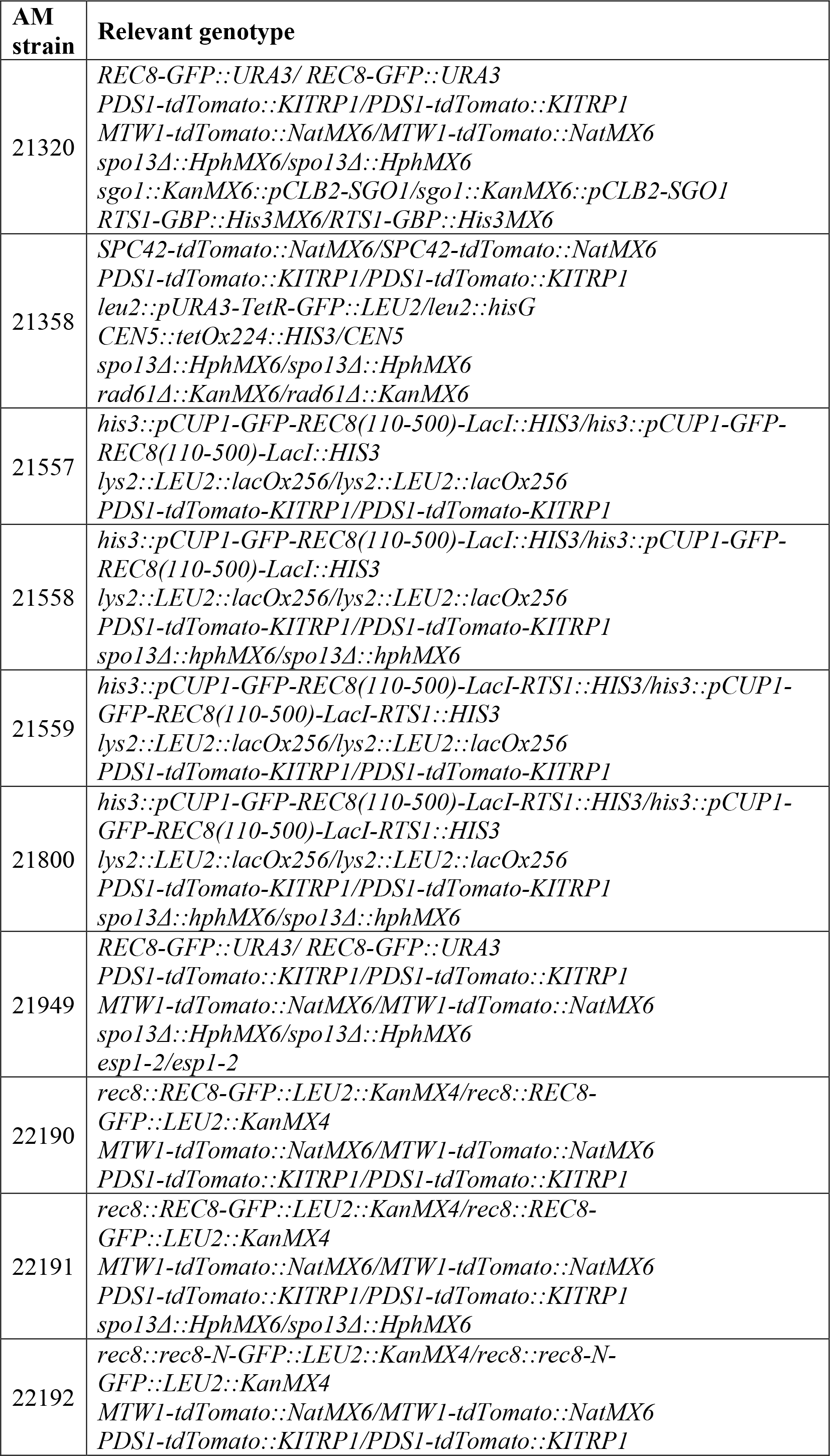
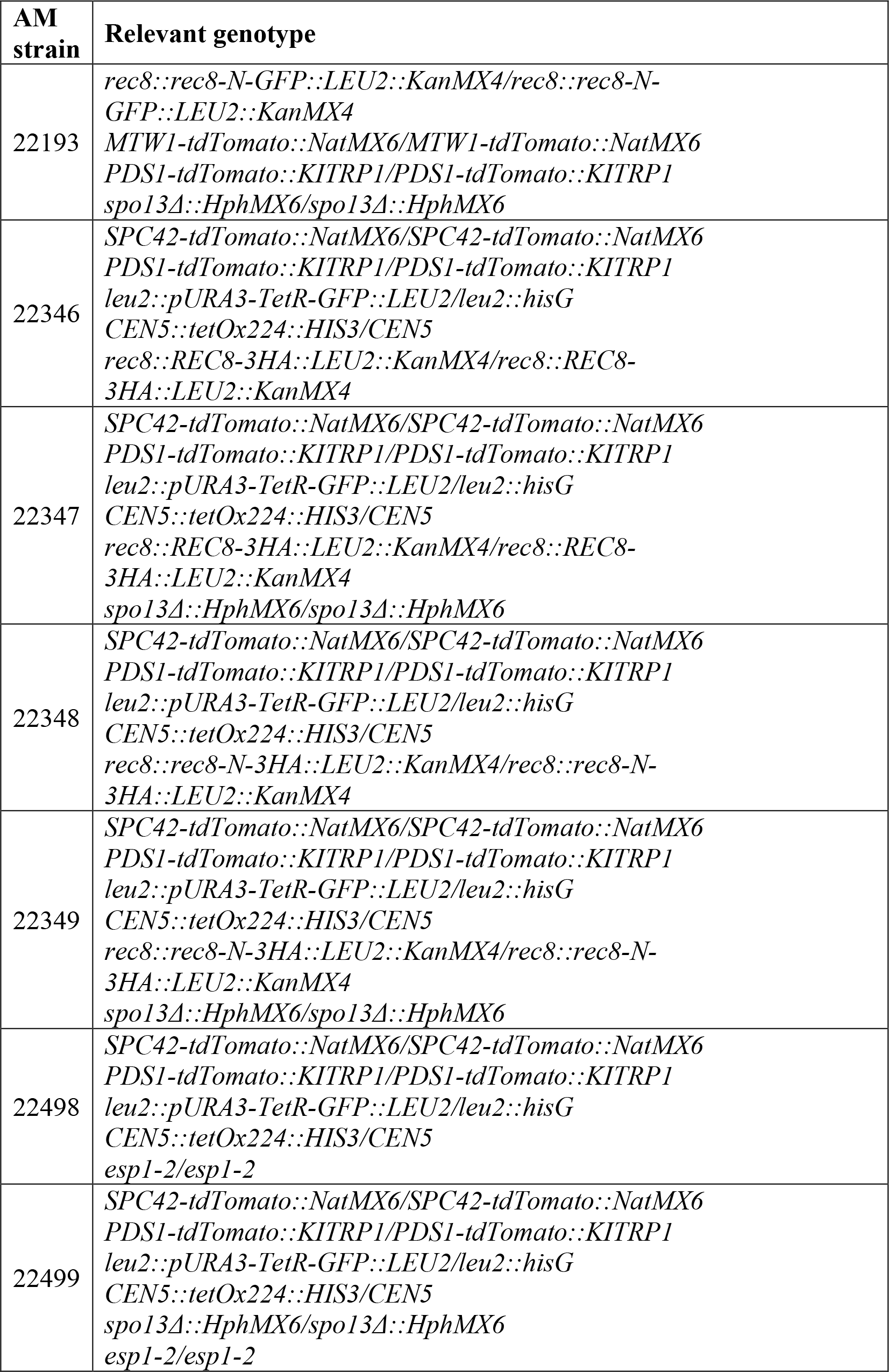
*Saccharomyces cerevisiae* strains used in this study.

| AM strain | Relevant genotype |
| --- | --- |
| 4015 | *ndt80Δ::LEU2/ndt80Δ::LEU2*<br>*REC8-3HA::URA3/ REC8-3HA::URA3* |
| 11633 | *ndt80Δ::LEU2/ndt80Δ::LEU2* |
| 13716 | *REC8-GFP::URA3/ REC8-GFP::URA3*<br>*PDS1-tdTomato::KITRP1/PDS1-tdTomato::KITRP1*<br>*MTW1-tdTomato::NatMX6/MTW1-tdTomato::NatMX6* |
| 15118 | *SPC42-tdTomato::NatMX6/SPC42-tdTomato::NatMX6*<br>*PDS1-tdTomato::KITRP1/PDS1-tdTomato::KITRP1*<br>*leu2::pURA3-TetR-GFP::LEU2/leu2::hisG*<br>*CEN5::tetOx224::HIS3/CEN5*<br>*spo13Δ::KanMX6/spo13Δ::KanMX6* |
| 15119 | *SPC42-tdTomato::NatMX6/SPC42-tdTomato::NatMX6*<br>*PDS1-tdTomato::KITRP1/PDS1-tdTomato::KITRP1*<br>*leu2::pURA3-TetR-GFP::LEU2/leu2::hisG*<br>*CEN5::tetOx224::HIS3/CEN5*<br>*mam1Δ::TRP1/mam1Δ::TRP1* |
| 15120 | *SPC42-tdTomato::NatMX6/SPC42-tdTomato::NatMX6*<br>*PDS1-tdTomato::KITRP1/PDS1-tdTomato::KITRP1*<br>*leu2::pURA3-TetR-GFP::LEU2/leu2::hisG*<br>*CEN5::tetOx224::HIS3/CEN5*<br>*spo13Δ::KanMX6/spo13Δ::KanMX6*<br>*mam1Δ::TRP1/mam1Δ::TRP1* |
| 15133 | *REC8-GFP::URA3/ REC8-GFP::URA3*<br>*PDS1-tdTomato::KITRP1/PDS1-tdTomato::KITRP1*<br>*MTW1-tdTomato::NatMX6/MTW1-tdTomato::NatMX6*<br>*spo13Δ::KanMX6/spo13Δ::KanMX6* |
| 15134 | *REC8-GFP::URA3/ REC8-GFP::URA3*<br>*PDS1-tdTomato::KITRP1/PDS1-tdTomato::KITRP1*<br>*MTW1-tdTomato::NatMX6/MTW1-tdTomato::NatMX6*<br>*mam1Δ::TRP1/mam1Δ::TRP1* |
| 15135 | *REC8-GFP::URA3/ REC8-GFP::URA3*<br>*PDS1-tdTomato::KITRP1/PDS1-tdTomato::KITRP1*<br>*MTW1-tdTomato::NatMX6/MTW1-tdTomato::NatMX6*<br>*spo13Δ::KanMX6/spo13Δ::KanMX6*<br>*mam1Δ::TRP1/mam1Δ::TRP1* |
| 15190 | *SPC42-tdTomato::NatMX6/SPC42-tdTomato::NatMX6*<br>*PDS1-tdTomato::KITRP1/PDS1-tdTomato::KITRP1*<br>*leu2::pURA3-TetR-GFP::LEU2/leu2::hisG*<br>*CEN5::tetOx224::HIS3/CEN5* |
| 15342 | *ndt80Δ::LEU2/ndt80Δ::LEU2*<br>*REC8-3HA::URA3/REC8-3HA::URA3*<br>*mam1Δ::TRP1/mam1Δ::TRP1* |
| 15343 | *ndt80Δ::LEU2/ndt80Δ::LEU2*<br>*REC8-3HA::URA3/REC8-3HA::URA3*<br>*spo13Δ::KanMX6/spo13Δ::KanMX6* |
| 15344 | *ndt80Δ::LEU2/ndt80Δ::LEU2*<br>*REC8-3HA::URA3/REC8-3HA::URA3*<br>*spo13Δ::KanMX6/spo13Δ::KanMX6*<br>*mam1Δ::TRP1/mam1Δ::TRP1* |
| 20033 | *REC8-GFP::URA3/ REC8-GFP::URA3*<br>*PDS1-tdTomato::KITRP1/PDS1-tdTomato::KITRP1*<br>*MTW1-tdTomato::NatMX6/MTW1-tdTomato::NatMX6*<br>*spo13Δ::HphMX6/spo13Δ::HphMX6* |
| 20868 | *REC8-GFP::URA3/ REC8-GFP::URA3*<br>*PDS1-tdTomato::KITRP1/PDS1-tdTomato::KITRP1*<br>*MTW1-tdTomato::NatMX6/MTW1-tdTomato::NatMX6*<br>*esp1-2/esp1-2* |
| 21068 | *SPC42-tdTomato::NatMX6/SPC42-tdTomato::NatMX6*<br>*PDS1-tdTomato::KITRP1/PDS1-tdTomato::KITRP1*<br>*leu2::pURA3-TetR-GFP::LEU2/leu2::hisG*<br>*CEN5::tetOx224::HIS3/CEN5*<br>*rad61Δ::KanMX6/rad61Δ::KanMX6* |
| 21315 | *REC8-GFP::URA3/ REC8-GFP::URA3*<br>*PDS1-tdTomato::KITRP1/PDS1-tdTomato::KITRP1*<br>*MTW1-tdTomato::NatMX6/MTW1-tdTomato::NatMX6*<br>*sgo1::KanMX6::pCLB2-SGO1/sgo1::KanMX6::pCLB2-SGO1* |
| 21316 | *REC8-GFP::URA3/ REC8-GFP::URA3*<br>*PDS1-tdTomato::KITRP1/PDS1-tdTomato::KITRP1*<br>*MTW1-tdTomato::NatMX6/MTW1-tdTomato::NatMX6*<br>*RTS1-GBP::His3MX6/RTS1-GBP::His3MX6* |
| 21317 | *REC8-GFP::URA3/ REC8-GFP::URA3*<br>*PDS1-tdTomato::KITRP1/PDS1-tdTomato::KITRP1*<br>*MTW1-tdTomato::NatMX6/MTW1-tdTomato::NatMX6*<br>*spo13Δ::HphMX6/spo13Δ::HphMX6*<br>*sgo1::KanMX6::pCLB2-SGO1/sgo1::KanMX6::pCLB2-SGO1* |
| 21318 | *REC8-GFP::URA3/ REC8-GFP::URA3*<br>*PDS1-tdTomato::KITRP1/PDS1-tdTomato::KITRP1*<br>*MTW1-tdTomato::NatMX6/MTW1-tdTomato::NatMX6*<br>*sgo1::KanMX6::pCLB2-SGO1/sgo1::KanMX6::pCLB2-SGO1*<br>*RTS1-GBP::His3MX6/RTS1-GBP::His3MX6* |
| 21319 | *REC8-GFP::URA3/ REC8-GFP::URA3*<br>*PDS1-tdTomato::KITRP1/PDS1-tdTomato::KITRP1*<br>*MTW1-tdTomato::NatMX6/MTW1-tdTomato::NatMX6*<br>*spo13Δ::HphMX6/spo13Δ::HphMX6*<br>*RTS1-GBP::His3MX6/RTS1-GBP::His3MX6* |
| 21320 | *REC8-GFP::URA3/ REC8-GFP::URA3*<br>*PDS1-tdTomato::KITRP1/PDS1-tdTomato::KITRP1*<br>*MTW1-tdTomato::NatMX6/MTW1-tdTomato::NatMX6*<br>*spo13Δ::HphMX6/spo13Δ::HphMX6*<br>*sgo1::KanMX6::pCLB2-SGO1/sgo1::KanMX6::pCLB2-SGO1*<br>*RTS1-GBP::His3MX6/RTS1-GBP::His3MX6* |
| 21358 | *SPC42-tdTomato::NatMX6/SPC42-tdTomato::NatMX6*<br>*PDS1-tdTomato::KITRP1/PDS1-tdTomato::KITRP1*<br>*leu2::pURA3-TetR-GFP::LEU2/leu2::hisG*<br>*CEN5::tetOx224::HIS3/CEN5*<br>*spo13Δ::HphMX6/spo13Δ::HphMX6*<br>*rad61Δ::KanMX6/rad61Δ::KanMX6* |
| 21557 | *his3::pCUP1-GFP-REC8(110-500)-LacI::HIS3/his3::pCUP1-GFP-REC8(110-500)-LacI::HIS3*<br>*lys2::LEU2::lacOx256/lys2::LEU2::lacOx256*<br>*PDS1-tdTomato-KITRP1/PDS1-tdTomato-KITRP1* |
| 21558 | *his3::pCUP1-GFP-REC8(110-500)-LacI::HIS3/his3::pCUP1-GFP-REC8(110-500)-LacI::HIS3*<br>*lys2::LEU2::lacOx256/lys2::LEU2::lacOx256*<br>*PDS1-tdTomato-KITRP1/PDS1-tdTomato-KITRP1*<br>*spo13Δ::hphMX6/spo13Δ::hphMX6* |
| 21559 | *his3::pCUP1-GFP-REC8(110-500)-LacI-RTS1::HIS3/his3::pCUP1-GFP-REC8(110-500)-LacI-RTS1::HIS3*<br>*lys2::LEU2::lacOx256/lys2::LEU2::lacOx256*<br>*PDS1-tdTomato-KITRP1/PDS1-tdTomato-KITRP1* |
| 21800 | *his3::pCUP1-GFP-REC8(110-500)-LacI-RTS1::HIS3/his3::pCUP1-GFP-REC8(110-500)-LacI-RTS1::HIS3*<br>*lys2::LEU2::lacOx256/lys2::LEU2::lacOx256*<br>*PDS1-tdTomato-KITRP1/PDS1-tdTomato-KITRP1*<br>*spo13Δ::hphMX6/spo13Δ::hphMX6* |
| 21949 | *REC8-GFP::URA3/ REC8-GFP::URA3*<br>*PDS1-tdTomato::KITRP1/PDS1-tdTomato::KITRP1*<br>*MTW1-tdTomato::NatMX6/MTW1-tdTomato::NatMX6*<br>*spo13Δ::HphMX6/spo13Δ::HphMX6*<br>*esp1-2/esp1-2* |
| 22190 | *rec8::REC8-GFP::LEU2::KanMX4/rec8::REC8-GFP::LEU2::KanMX4*<br>*MTW1-tdTomato::NatMX6/MTW1-tdTomato::NatMX6*<br>*PDS1-tdTomato::KITRP1/PDS1-tdTomato::KITRP1* |
| 22191 | *rec8::REC8-GFP::LEU2::KanMX4/rec8::REC8-GFP::LEU2::KanMX4*<br>*MTW1-tdTomato::NatMX6/MTW1-tdTomato::NatMX6*<br>*PDS1-tdTomato::KITRP1/PDS1-tdTomato::KITRP1*<br>*spo13Δ::HphMX6/spo13Δ::HphMX6* |
| 22192 | *rec8::rec8-N-GFP::LEU2::KanMX4/rec8::rec8-N-GFP::LEU2::KanMX4*<br>*MTW1-tdTomato::NatMX6/MTW1-tdTomato::NatMX6*<br>*PDS1-tdTomato::KITRP1/PDS1-tdTomato::KITRP1* |
| 22193 | *rec8::rec8-N-GFP::LEU2::KanMX4/rec8::rec8-N-GFP::LEU2::KanMX4*<br>*MTW1-tdTomato::NatMX6/MTW1-tdTomato::NatMX6*<br>*PDS1-tdTomato::KITRP1/PDS1-tdTomato::KITRP1*<br>*spo13Δ::HphMX6/spo13Δ::HphMX6* |
| 22346 | *SPC42-tdTomato::NatMX6/SPC42-tdTomato::NatMX6*<br>*PDS1-tdTomato::KITRP1/PDS1-tdTomato::KITRP1*<br>*leu2::pURA3-TetR-GFP::LEU2/leu2::hisG*<br>*CEN5::tetOx224::HIS3/CEN5*<br>*rec8::REC8-3HA::LEU2::KanMX4/rec8::REC8-3HA::LEU2::KanMX4* |
| 22347 | *SPC42-tdTomato::NatMX6/SPC42-tdTomato::NatMX6*<br>*PDS1-tdTomato::KITRP1/PDS1-tdTomato::KITRP1*<br>*leu2::pURA3-TetR-GFP::LEU2/leu2::hisG*<br>*CEN5::tetOx224::HIS3/CEN5*<br>*rec8::REC8-3HA::LEU2::KanMX4/rec8::REC8-3HA::LEU2::KanMX4*<br>*spo13Δ::HphMX6/spo13Δ::HphMX6* |
| 22348 | *SPC42-tdTomato::NatMX6/SPC42-tdTomato::NatMX6*<br>*PDS1-tdTomato::KITRP1/PDS1-tdTomato::KITRP1*<br>*leu2::pURA3-TetR-GFP::LEU2/leu2::hisG*<br>*CEN5::tetOx224::HIS3/CEN5*<br>*rec8::rec8-N-3HA::LEU2::KanMX4/rec8::rec8-N-3HA::LEU2::KanMX4* |
| 22349 | *SPC42-tdTomato::NatMX6/SPC42-tdTomato::NatMX6*<br>*PDS1-tdTomato::KITRP1/PDS1-tdTomato::KITRP1*<br>*leu2::pURA3-TetR-GFP::LEU2/leu2::hisG*<br>*CEN5::tetOx224::HIS3/CEN5*<br>*rec8::rec8-N-3HA::LEU2::KanMX4/rec8::rec8-N-3HA::LEU2::KanMX4*<br>*spo13Δ::HphMX6/spo13Δ::HphMX6* |
| 22498 | *SPC42-tdTomato::NatMX6/SPC42-tdTomato::NatMX6*<br>*PDS1-tdTomato::KITRP1/PDS1-tdTomato::KITRP1*<br>*leu2::pURA3-TetR-GFP::LEU2/leu2::hisG*<br>*CEN5::tetOx224::HIS3/CEN5*<br>*esp1-2/esp1-2* |
| 22499 | *SPC42-tdTomato::NatMX6/SPC42-tdTomato::NatMX6*<br>*PDS1-tdTomato::KITRP1/PDS1-tdTomato::KITRP1*<br>*leu2::pURA3-TetR-GFP::LEU2/leu2::hisG*<br>*CEN5::tetOx224::HIS3/CEN5*<br>*spo13Δ::HphMX6/spo13Δ::HphMX6*<br>*esp1-2/esp1-2* |

**Table S2.** Plasmids generated in this study

| Plasmid | Description | Purpose and notes |
| --- | --- | --- |
| AMp1312 | pFA6a-HphMX6-pCLB2-3HA | Plasmid for promoter replacement with *pCLB2*. |
| AMp1317 | YIplac128-REC8-GFP | *LEU2* integration plasmid carrying *REC8-GFP*. |
| AMp1368 | YIplac128-rec8-N-GFP | *LEU2* integration plasmid carrying *rec8-N-GFP*. |

**Table S3.**
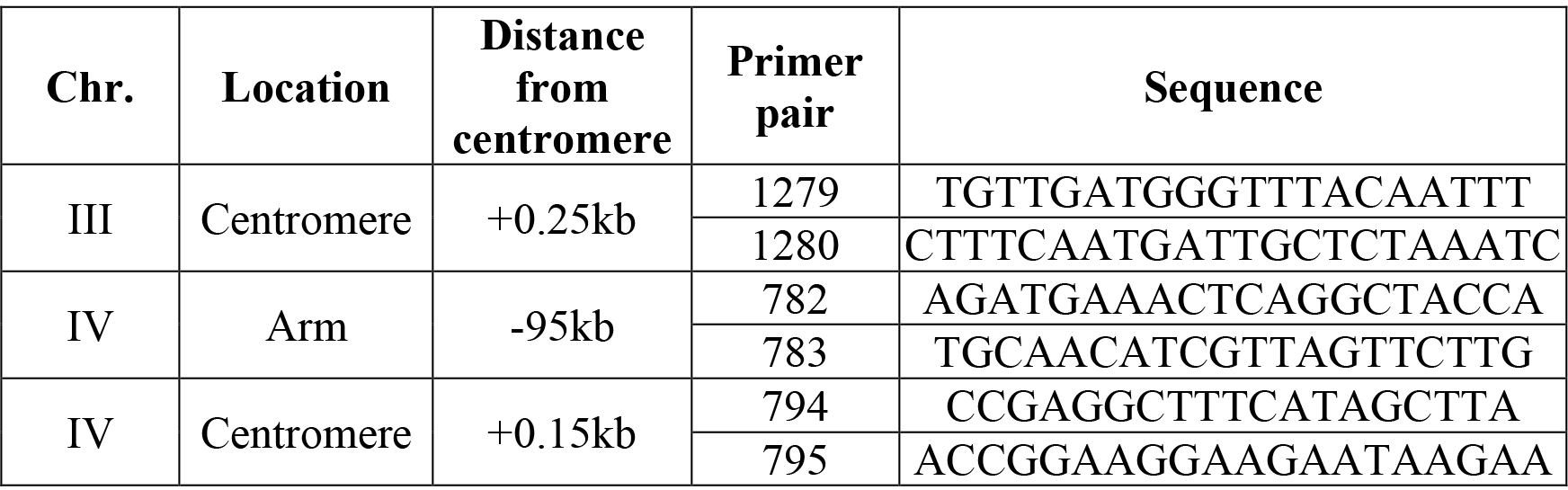
qPCR primers used in this study. For distances from centromeres, “−” indicates the location is upstream of the centromere, whereas “+” indicates the location is downstream of the centromere.

## References

Argüello-Miranda O, Zagoriy I, Mengoli V, Rojas J, Jonak K, Oz T, Graf P,Zachariae W. 2017. Casein Kinase 1 Coordinates Cohesin Cleavage, Gametogenesis, and Exit from M Phase in Meiosis II. Dev Cell 40: 37–52.

Attner MA, Miller MP, Ee L-S, Elkin SK, Amon A. 2013. Polo kinase Cdc5 is a central regulator of meiosis I. Proc Natl Acad Sci USA 110: 14278–14283.

Ben-Shahar TR, Heeger S, Lehane C, East P, Flynn H, Skehel M, Uhlmann F. 2008. Eco1-Dependent Cohesin Acetylation During Establishment of Sister Chromatid Cohesion. Science 321: 563–566.

Brar GA, Kiburz BM, Zhang Y, Kim J-E, White F, Amon A. 2006. Rec8 phosphorylation and recombination promote the step-wise loss of cohesins in meiosis. Nature 441: 532–536.

Buheitel J, Stemmann O. 2013. Prophase pathway-dependent removal of cohesin from human chromosomes requires opening of the Smc3-Scc1 gate. EMBO J 32: 666–676.

Buonomo SB, Clyne RK, Fuchs J, Loidl J, Uhlmann F, Nasmyth K. 2000. Disjunction of homologous chromosomes in meiosis I depends on proteolytic cleavage of the meiotic cohesin Rec8 by separin. Cell 103: 387–398.

Challa K, Lee M-S, Shinohara M, Kim KP, Shinohara A. 2016. Rad61/Wpl1 (Wapl), a cohesin regulator, controls chromosome compaction during meiosis. Nucleic Acids Res gkw034.

Challa K, Shinohara M, Klein F, Gasser SM, Shinohara A. 2018. Meiosis-specific prophase-like pathway controls cleavage-independent release of cohesin by Wapl. bioRxiv 250589.

Fox C, Zou J, Rappsilber J, Marston AL. 2017. Cdc14 phosphatase directs centrosome re-duplication at the meiosis I to meiosis II transition in budding yeast. Wellcome Open Res 2: 2.

Gauss R, Trautwein M, Sommer T, Spang A. 2005. New modules for the repeated internal and N-terminal epitope tagging of genes in Saccharomyces cerevisiae. Yeast 22: 1–12.

Gruber S, Haering CH, Nasmyth K. 2003. Chromosomal cohesin forms a ring. Cell 112: 765–777.

Haering CH, Löwe J, Hochwagen A, Nasmyth K. 2002. Molecular architecture of SMC proteins and the yeast cohesin complex. Mol Cell 9: 773–788.

Hartman T, Stead K, Koshland D, Guacci V. 2000. Pds5p is an essential chromosomal protein required for both sister chromatid cohesion and condensation in Saccharomyces cerevisiae. J Cell Biol 151: 613–626.

Hassold T, Hunt P. 2001. To err (meiotically) is human: the genesis of human aneuploidy. Nat Rev Genet 2: 280–291.

Jonak K, Zagoriy I, Oz T, Graf P, Rojas J, Mengoli V, Zachariae W. 2017. APC/C-Cdc20 mediates deprotection of centromeric cohesin at meiosis II in yeast. Cell Cycle 16: 1145–1152.

Katis VL, Gálová M, Rabitsch KP, Gregan J, Nasmyth K. 2004a. Maintenance of cohesin at centromeres after meiosis I in budding yeast requires a kinetochore-associated protein related to MEI-S332. Curr Biol 14: 560–572.

Katis VL, Lipp JJ, Imre R, Bogdanova A, Okaz E, Habermann B, Mechtler K, Nasmyth K, Zachariae W. 2010. Rec8 phosphorylation by casein kinase 1 and Cdc7-Dbf4 kinase regulates cohesin cleavage by separase during meiosis. Supplement. Dev Cell 18: 397–409.

Katis VL, Matos J, Mori S, Shirahige K, Zachariae W, Nasmyth K. 2004b. Spo13 facilitates monopolin recruitment to kinetochores and regulates maintenance of centromeric cohesion during yeast meiosis. Curr Biol 14: 2183–2196.

Kiburz BM, Reynolds DB, Megee PC, Marston AL, Lee B, Lee TI, Levine SS, Young RA, Amon A. 2005. The core centromere and Sgo1 establish a 50-kb cohesin-protected domain around centromeres during meiosis I. Genes Dev 19: 3017–3030.

Kim J, Ishiguro K-I, Nambu A, Akiyoshi B, Yokobayashi S, Kagami A, Ishiguro T, Pendás AM, Takeda N, Sakakibara Y, et al.. 2014. Meikin is a conserved regulator of meiosis-I-specific kinetochore function. Nature 517: 466–471.

Kitajima TS, Kawashima SA, Watanabe Y. 2004. The conserved kinetochore protein shugoshin protects centromeric cohesion during meiosis. Nature 427: 510–517.

Kitajima TS, Sakuno T, Ishiguro K-I, Iemura S-I, Natsume T, Kawashima SA, Watanabe Y. 2006. Shugoshin collaborates with protein phosphatase 2A to protect cohesin. Nature 441: 46–52.

Klapholz S, Esposito RE. 1980. Recombination and chromosome segregation during the single division meiosis in SPO12-1 and SPO13-1 diploids. Genetics 96: 589–611.

Klein F, Mahr P, Gálová M, Buonomo SB, Michaelis C, Nairz K, Nasmyth K. 1999. A central role for cohesins in sister chromatid cohesion, formation of axial elements, and recombination during yeast meiosis. Cell 98: 91–103.

Knop M, Siegers K, Pereira G, Zachariae W, Winsor B, Nasmyth K, Schiebel E.1999. Epitope tagging of yeast genes using a PCR-based strategy: more tags and improved practical routines. Yeast 15: 963–972.

Lafont AL, Song J, Rankin S. 2010. Sororin cooperates with the acetyltransferase Eco2 to ensure DNA replication-dependent sister chromatid cohesion. Proc Natl Acad Sci USA 107: 20364–20369.

Lee B, Amon A. 2003. Role of Polo-like kinase CDC5 in programming meiosis I chromosome segregation. Science 300: 482–486.

Lee B, Kiburz BM, Amon A. 2004. Spo13 maintains centromeric cohesion and kinetochore coorientation during meiosis I. Curr Biol 14: 2168–2182.

Lee J, Kitajima TS, Tanno Y, Yoshida K, Morita T, Miyano T, Miyake M, Watanabe Y. 2008. Unified mode of centromeric protection by shugoshin in mammalian oocytes and somatic cells. Nat Cell Biol 10: 42–52.

Liu H, Jia L, Yu H. 2013a. Phospho-H2A and Cohesin Specify Distinct Tension-Regulated Sgo1 Pools at Kinetochores and Inner Centromeres. Curr Biol.

Liu H, Rankin S, Yu H. 2013b. Phosphorylation-enabled binding of SGO1-PP2A to cohesin protects sororin and centromeric cohesion during mitosis. Nat Cell Biol 15: 40–49.

Longtine MS, McKenzie A, Demarini DJ, Shah NG, Wach A, Brachat A, Philippsen P, Pringle JR. 1998. Additional modules for versatile and economical PCR-based gene deletion and modification in Saccharomyces cerevisiae. Yeast 14: 953–961.

Losada A, Hirano M, Hirano T. 1998. Identification of Xenopus SMC protein complexes required for sister chromatid cohesion. Genes Dev 12: 1986–1997.

Marston AL, Amon A. 2004. Meiosis: cell-cycle controls shuffle and deal. Nat Rev Mol Cell Biol 5: 983–997.

Marston AL, Tham W-H, Shah H, Amon A. 2004. A genome-wide screen identifies genes required for centromeric cohesion. Science 303: 1367–1370.

Matos J, Lipp JJ, Bogdanova A, Guillot S, Okaz E, Junqueira M, Shevchenko A, Zachariae W. 2008. Dbf4-dependent CDC7 kinase links DNA replication to the segregation of homologous chromosomes in meiosis I. Cell 135: 662–678.

Michaelis C, Ciosk R, Nasmyth K. 1997. Cohesins: chromosomal proteins that prevent premature separation of sister chromatids. Cell 91: 35–45.

Miyazaki S, Kim J, Yamagishi Y, Ishiguro T, Okada Y, Tanno Y, Sakuno T, Watanabe Y. 2017. Meikin-associated polo-like kinase specifies Bub1 distribution in meiosis I. Genes Cells 22: 552–567.

Moqtaderi Z, Struhl K. 2008. Expanding the repertoire of plasmids for PCR-mediated epitope tagging in yeast. Yeast 25: 287–292.

Nerusheva OO, Galander S, Fernius J, Kelly D, Marston AL. 2014. Tension-dependent removal of pericentromeric shugoshin is an indicator of sister chromosome biorientation. Genes Dev 28: 1291–1309. http://genesdev.cshlp.org/content/28/12/1291.full.

Nishiyama T, Ladurner R, Schmitz J, Kreidl E, Schleiffer A, Bhaskara V, Bando M, Shirahige K, Hyman AA, Mechtler K, et al.. 2010. Sororin mediates sister chromatid cohesion by antagonizing Wapl. Cell 143: 737–749.

Panizza S, Tanaka T, Hochwagen A, Eisenhaber F, Nasmyth K. 2000. Pds5 cooperates with cohesin in maintaining sister chromatid cohesion. Curr Biol 10: 1557–1564.

Rankin S, Ayad NG, Kirschner MW. 2005. Sororin, a substrate of the anaphase-promoting complex, is required for sister chromatid cohesion in vertebrates. Mol Cell 18: 185–200.

Riedel CG, Katis VL, Katou Y, Mori S, Itoh T, Helmhart W, Gálová M, Petronczki M, Gregan J, Cetin B, et al.. 2006. Protein phosphatase 2A protects centromeric sister chromatid cohesion during meiosis I. Nature 441: 53–61.

Rothbauer U, Zolghadr K, Tillib S, Nowak D, Schermelleh L, Gahl A, Backmann N, Conrath K, Muyldermans S, Cardoso MC, et al.. 2006. Targeting and tracing antigens in live cells with fluorescent nanobodies. Nat Methods 3: 887–889. http://www.cardoso-lab.org/publications/Rothbauer_2006.pdf.

Salah SM, Nasmyth K. 2000. Destruction of the securin Pds1p occurs at the onset of anaphase during both meiotic divisions in yeast. Chromosoma 109: 27–34. https://link.springer.com/article/10.1007%2Fs004120050409.

Schmitz J, Watrin E, Lénárt P, Mechtler K, Peters J-M. 2007. Sororin is required for stable binding of cohesin to chromatin and for sister chromatid cohesion in interphase. Curr Biol 17: 630–636.

Shonn MA, McCarroll R, Murray AW. 2002. Spo13 protects meiotic cohesin at centromeres in meiosis I. Genes Dev 16: 1659–1671.

Sumara I, Vorlaufer E, Gieffers C, Peters BH, Peters JM. 2000. Characterization of vertebrate cohesin complexes and their regulation in prophase. J Cell Biol 151: 749–762.

Tang Z, Shu H, Qi W, Mahmood NA, Mumby MC, Yu H. 2006. PP2A is required for centromeric localization of Sgo1 and proper chromosome segregation. Dev Cell 10: 575–585.

Tóth A, Ciosk R, Uhlmann F, Gálová M, Schleiffer A, Nasmyth K. 1999. Yeast Cohesin complex requires a conserved protein, Eco1p(Ctf7), to establish cohesion between sister chromatids during DNA replication. Genes Dev 13: 320–333.

Tóth A, Rabitsch KP, Gálová M, Schleiffer A, Buonomo SB, Nasmyth K. 2000. Functional genomics identifies monopolin: a kinetochore protein required for segregation of homologs during meiosis I. Cell 103: 1155–1168.

Unal E, Heidinger-Pauli JM, Kim W, Guacci V, Onn I, Gygi SP, Koshland DE. 2008. A molecular determinant for the establishment of sister chromatid cohesion. Science 321: 566–569.

Vincenten N, Kuhl L-M, Lam I, Oke A, Kerr AR, Hochwagen A, Fung J, Keeney S, Vader G, Marston AL. 2015. The kinetochore prevents centromere-proximal crossover recombination during meiosis. elife 4: 923.

Waizenegger IC, Hauf S, Meinke A, Peters JM. 2000. Two distinct pathways remove mammalian cohesin from chromosome arms in prophase and from centromeres in anaphase. Cell 103: 399–410.

Wang HT, Frackman S, Kowalisyn J, Esposito RE, Elder R. 1987. Developmental regulation of SPO13, a gene required for separation of homologous chromosomes at meiosis I. Mol Cell Biol 7: 1425–1435.

Warren WD, Steffensen S, Lin E, Coelho P, Loupart M, Cobbe N, Lee JY, McKay MJ, Orr-Weaver T, Heck MM, et al.. 2000. The Drosophila RAD21 cohesin persists at the centromere region in mitosis. Curr Biol 10: 1463–1466.

Watanabe Y, Nurse P. 1999. Cohesin Rec8 is required for reductional chromosome segregation at meiosis. Nature 400: 461–464.

Yaakov G, Thorn K, Morgan DO. 2012. Separase Biosensor Reveals that Cohesin Cleavage Timing Depends on Phosphatase PP2A(Cdc55) Regulation. Dev Cell 23: 124–136.

Yu H-G, Koshland D. 2005. Chromosome morphogenesis: condensin-dependent cohesin removal during meiosis. Cell 123: 397–407.

